# *InSegtCone*: Interactive Segmentation of crystalline Cones in compound eyes

**DOI:** 10.1101/2020.12.15.422850

**Authors:** Pierre Tichit, Tunhe Zhou, Hans Martin Kjer, Vedrana Andersen Dahl, Anders Bjorholm Dahl, Emily Baird

## Abstract

Understanding the diversity of eyes is crucial to unravel how different animals use vision to interact with their respective environments. To date, comparative studies of eye anatomy are scarce because they often involve time-consuming or inefficient methods. X-ray micro-tomography is a promising high-throughput imaging technique that enables to reconstruct the 3D anatomy of eyes, but powerful tools are needed to perform fast conversions of anatomical reconstructions into functional eye models. We developed a computing method named *InSegtCone* to automatically segment the crystalline cones in the apposition compound eyes of arthropods. Here, we describe the full auto-segmentation process, showcase its application to three different insect compound eyes and evaluate its performance. The auto-segmentation could successfully label the full individual shapes of 60%-80% of the crystalline cones, and is about as accurate and 250 times faster than manual labelling of the individual cones. We believe that *InSegtCone* can be an important tool for peer scientists to enable extensive comparisons of the diversity of eyes and vision in arthropods.

## Introduction

Arthropods comprise more than 80% of animals living on the planet (Roskov et al., 2000) but little is known about the diversity of the visual systems in this phylum (Rosenthal et al. 2017). Many arthropods make extensive use of visual information to perform essential behaviours, such as locating suitable mates, finding food or avoiding predators (Cronin et al., 2014). To better understand how these animals interact with their environment, it is therefore important to carry out anatomical and functional investigations of the myriad of arthropod eyes. Moreover, better understanding the eyes of arthropods will not only inform us about the visual ecology of these species, but also about fundamental aspects of animal vision.

Most Arthropods have a pair of compound eyes consisting of repeated units called ommatidia. An ommatidium typically consists of three elements: an external lens that forms a regular facet visible on the external surface of the eye, a crystalline cone and a rhabdom (Land and Nilsson, 2012). The light-sensitive rhabdom is the ‘sensor’ unit that collects light provided by unique (in apposition compound eyes) or multiple (in superposition compound eyes) transparent ‘optics’ unit – the lens and the crystalline cone. Each ommatidium samples light from a small angular portion of the world that, once integrated, enables arthropods to generate detailed and colourful images across the field of view (FOV) of the compound eye.

To be able to resolve an object in a given light regime, each portion of a compound eye trades-off adequate spatial resolution and sufficient light capture, or optical sensitivity (Land, 1997). This is because spatial resolution is ultimately determined by the angular spacing of the neighbouring ommatidia – the inter-ommatidial (IO) angle – such that a reduction in IO angle increases resolution. On the contrary, optical sensitivity depends on the angular area over which photons are captured; a bigger IO angle therefore leads to better sensitivity. The optical properties of ommatidia thus typically vary across the topology of compound eyes to optimise resolution and/or sensitivity in different parts of the FOV in a way that reflects the ecology of the animal. For example, the frontal region of the eye of the male carpenter bee *Xylocopa tenuiscapa* has enlarged ommatidia with small IO angle, which enhances both sensitivity and resolution and probably is an adaptation for detecting and chasing mates (Somanathan et al., 2017). Thus, anatomical comparisons of compound eyes across sexes, castes, life-stages, populations or species, provide formidable opportunities to better understand the visual ecology and behaviour of arthropods.

Exploring how the ecology of arthropods relates to their visual anatomy requires large-scale and detailed analysis of eye structures, something that is challenging for two major reasons. Firstly, it necessitates investigations into the inner anatomy of compound eyes, because properties such as the elongation axes of crystalline cones and the diameters and lengths of rhabdoms are needed for accurate calculations of optical sensitivity and IO angle (Land, 1997). Furthermore, these properties must be measured in high-resolution across the eye to obtain reliable topological information (Taylor et al., 2019a). Traditional methods to study visual anatomy provide incomplete information and/or are too time consuming to enable large-scale analyses. Imaging methods such as Transmission Electron Microscopy (TEM), a technique that is typically used to measure rhabdom properties, are slow and produce coarse topologies from only a few slices in each eye sample. The pseudopupil technique (Berry et al., 2007), which measures the topology of IO angle, is relatively fast but requires live animals and typically generates coarse resolution maps that do not include the full extent of the FOV (Taylor et al., 2019a), sometimes filtering out precious fine-scale information (Bagheri et al., 2020).

How can we quickly obtain fine-scale topological data, both on the outside and the inside of compound eyes that typically possess thousands of ommatidia? X-ray microtomography (micro-CT) is a promising method to generate fast (a scan typically lasts from a few minutes to a few hours) and accurate (with isotropic spatial resolution of a micrometre or less) 3D models of an eye (Baird and Taylor, 2017). This method has the advantage of keeping the eye geometry intact, unlike 2D methods that inherently lose part of the geometrical information through the physical sectioning process. In a recent study, Taylor et al. (2019a) used micro-CT to obtain maps of the optical properties across the entire FOV of *Bombus terrestris* compound eyes in unprecedented detail to explore the effect of body size on the topological scaling of visual parameters.

Paradoxically, micro-CT generates more information than is currently feasible to process, in particular, it provides a high level of detail about the geometry of the rhabdoms and crystalline cones that has been left out of previous modelling approaches (e.g. Taylor et al., 2019a, 2020). Information about the geometry of individual rhabdoms would enable optical sensitivity calculations, and measuring the orientation of crystalline cones is crucial to produce accurate estimates of the visual IO angle over the entire eye surface (Stavenga, 1979). This is because crystalline cones are often skewed relative to the surface of the cornea, which is visible in *Figure 1d* in Taylor et al. (2019a), so that the topology of their viewing directions determines the angular spacing of neighbouring ommatidia (Baumgärtner, 1928; Stavenga, 1979). In other words, measurements solely based on the angular spacing of facets on the eye surface, called the corneal IO angle, generate biased estimates of resolution and FOV (Bergman and Rutowski, 2016; Taylor et al., 2019a). This difference is particularly striking at the edges of the FOV, where corneal measurements of the IO angle often overestimate the actual visual IO angle (Seidl and Kaiser, 1981; Taylor et al., 2019a). To address this problem, manual segmentation (or labelling) of each crystalline cone ‘by hand’ in volumetric analysis programs is an option. Unfortunately, unless it is possible to infer the elongation axes of the cones from their specific geometry, as is the case in fiddler crabs (Bagheri et al., 2020), manual segmentation across more than a few samples becomes nearly unfeasible for eyes that frequently consist of several thousands of ommatidia. More generally, the current time-limiting factor of micro-CT is often not the scanning itself but the volumetric analysis of the images it produces, particularly when segmentation is required. If it is to become a widespread tool for large-scale anatomical studies in arthropod vision, new methods are needed to automate the analysis of micro-CT scans.

**Figure 1:**
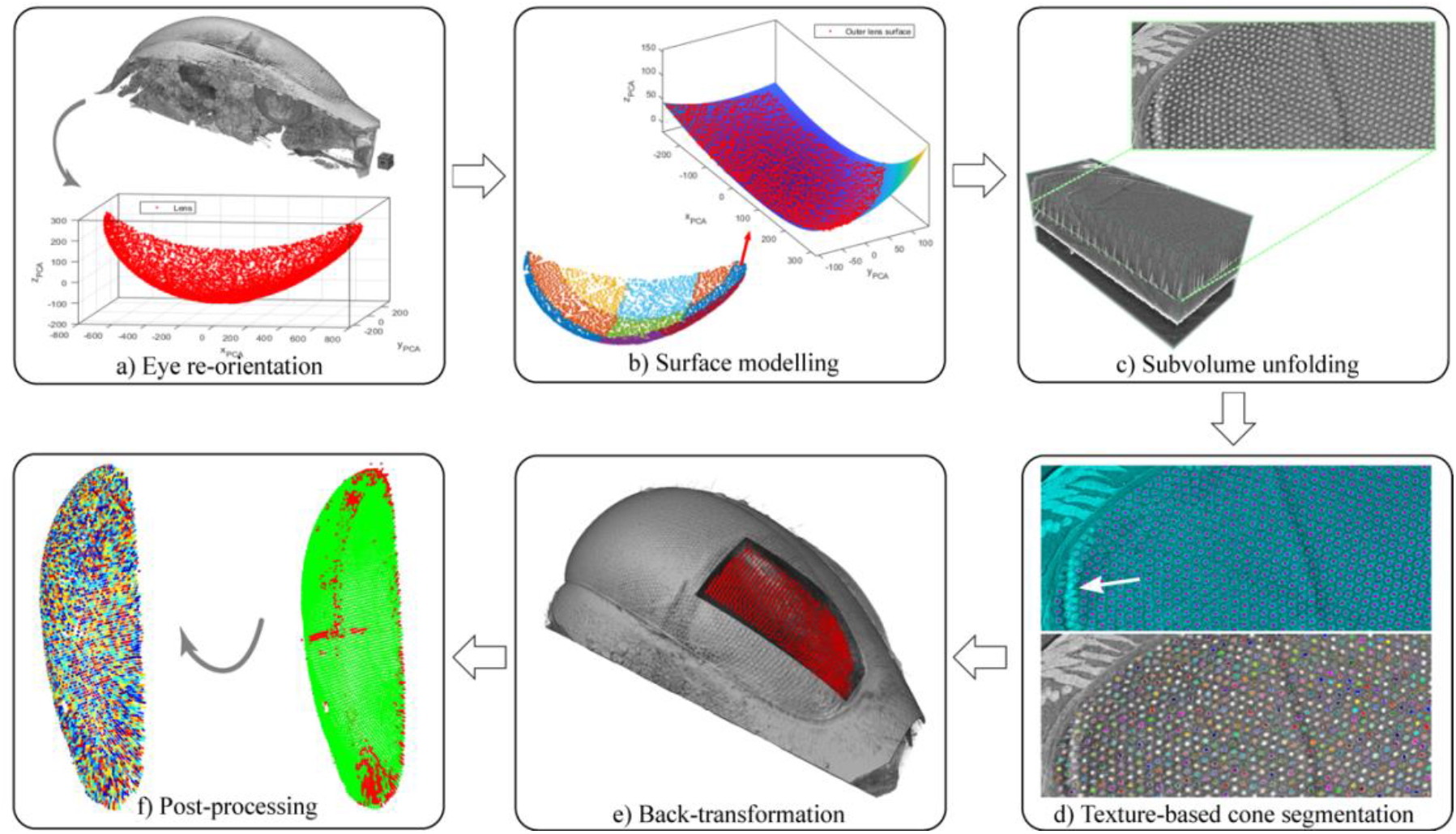
Outline of the process to automatically segment crystalline cones. The method is illustrated with images of *Apis mellifera* as an example. *(a)* Reorientation of the eye (volume rendering) and the labelled external cornea (red) into the ‘PCA space’. *(b)* Division of the cornea surface voxels into subregions (each plotted with a different colour) and modelling of each subregion using polynomial fitting in its ‘PCA space’ (top). *(c)* Extraction and unfolding of a subvolume of the eye data. Note that the cornea surface in the unfolded subvolume is almost flat, so that the pattern of crystalline cones in the cross-section is regular. *(d)* Auto-segmentation of the raw cones using a texture-based approach. The results of this step are displayed on one slice from the unfolded images (top) and in the whole eye subvolume (bottom). The white arrow indicates the location of a circular artefact that locally prevented the segmentation of a few cones. *(e)* Back-transformation of the raw auto-segmented cones into the original eye volume (3D rendering). *(f)* Post-processing of the raw cones labels (right subfigure) to eliminate noisy detections (red on the right subfigure) and retain valid cones (green on the right subfigure - left subfigure).

In this paper, we start to bridge this gap by developing a computing method, *InSegtCone*, that segments crystalline cones in arthropod compound eyes with little input from the user, i.e. nearly automatically. Our auto-segmentation method is based on an algorithm for interactive segmentation (Dahl et al., 2020) that automatically labels repeated objects after a short manual training. Here, we explore the functionality of *InSegtCone* by applying it to three insect species with differently shaped apposition compound eyes: the Western honeybee *Apis mellifera*, the buff-tailed bumblebee *Bombus terrestris*, and the green-veined white butterfly *Pieris napi*. We demonstrate that the auto-segmentation process can successfully extract the full shapes of 60%-80% of the total number of crystalline cones (~6000). We evaluate the performance of the *InSegtCone* and compare it to manual labelling. We then discuss the remaining limitations and new opportunities that this new technique generates.

## Material and Methods

### Study animals

Micro-CT image stacks of eye samples of adult workers (females) of *Apis mellifera* (specimen: LU:3_14:AM_F_5) and *Bombus terrestris* (specimen: LU:4_16_: BT_F_CE_10) were the same as those previously analysed in Taylor et al. (2019b), where the sample preparation and data acquisition method are described in detail. An adult female of *Pieris napi* was obtained from laboratory stock at the Department of Zoology, Stockholm University, Sweden.

### Sample preparation

The left or right compound eyes of the bee specimens were dissected, fixated, stained, and embedded in epoxy resin according to the procedure described in Taylor et al. (2019a). The left half head of the sample of *P. napi* was fixated for 7 days at 4°C in a 0.5% phosphotungstic acid (PTA) solution (0.5 mg/mL of PTA in 70/30% ethanol/water solution) for staining. The sample of *P. napi* was not embedded in resin but scanned directly in a 70% ethanol solution.

### X-ray microtomography (micro-CT)

Micro-CT imaging of bee eyes was conducted at *Diamond Light Source* Beamline I13-2 (Peić et al., 2013; Rau et al., 2011), Harwell Science and Innovation Campus, Oxfordshire (UK). The voxel size of the bee eyes was 1.6 μm. A detailed description of the scanning procedure can be found in Taylor et al. (2019a).

The *P. napi* sample was scanned at the *Stockholm Brain Imaging Center – SUBIC* (Lacerda and Lindblom, 2014) at Stockholm University (Sweden) using the 3D submicron imaging system Xradia Versa 520 (Zeiss, Jena, Germany). The imaging was performed with the X-ray source running with a voltage of 80 kV and a power of 7 W. A 4x optical objective was used, combined with the geometrical magnification, leading to an effective voxel size of 1.08 μm. The tomography contained 2401 projections over 360°. The exposure time was 1 s per projection. The projection images were then reconstructed automatically using the Zeiss Scout-and-Scan software.

### Volumetric segmentation of the eye and the external cornea

To perform the auto-segmentation of the cones, *InSegtCone* required input labels of key eye features that were extracted from the reconstructed microCT images. The external cornea and the full eye volume of each compound eye were labelled by volumetric segmentation in Amira (FEI, Hillsboro, USA) using a method modified from Taylor et al. (2019a) . The original 32-bit images reconstructed from the scans were cropped and re-saved in 8-bit files in the program Drishti Paint (Limaye, 2012). The images were resampled to 4 μm voxels. The eye volume was labelled thanks to a combination of automatic thresholding, filling of holes on each slice, shrinking and growing of the labelled volume, and selection of connected components. The outer surface of the cornea was extracted from the eye label by manually drawing a geodesic path at the border of the cornea and extracting the enclosed surface. The two labels were saved as volumetric images for later analysis in MATLAB (The MathWorks Inc., USA).

### Manual segmentation of the crystalline cones

Crystalline cones were labelled manually in the full-resolution 8-bit images using the brush tool of Amira across 2D slices. Care was taken to obtain a collection of segmented cones distributed as uniformly as possible across the eye. To compare with the segmentation time of the automatic method, the time required for a user to perform manual segmentation of the cones of *P. napi* in Amira was estimated.

### Automatic segmentation of the crystalline cones

The schematic of the automatic segmentation (or auto-segmentation) process of the crystalline cones called *InSegtCone* is described in *Figure 1*. The process was performed in MATLAB and the code is available for download on *Github* (link will be provided before publishing). The main innovative aspect of the segmentation method consists in the unfolding of the original eye volume into a 2D coordinate system referenced to the external surface of the cornea. This simplifies the segmentation problem, as the actual texture-based segmentation (Dahl et al., 2020) can then take place in two instead of three dimensions. To allow accurate unfolding, despite the variable curvature of the external surface of the cornea, the eye was divided into sub-regions prior to the unfolding. The effect of this step on the performance of the method was tested by independently applying the auto-segmentation method to the eye of *A. mellifera* divided into increasing numbers of sub-regions (1, 2, 4, 6, 9, 12). The eye of *B. terrestris* (resp. *P.napi*) was divided into 12 (resp. 9) sub-regions.

The crystalline cone auto-segmentation process consisted of six steps: *(a)* reorientation of the eye; *(b)* division of the outer cornea voxels into sub-regions and surface modelling using polynomial fitting; *(c)* extraction and unfolding of a sub-volume of the data capturing the crystalline cone layer; *(d)* auto-segmentation of the cones using a texture-based approach; *(e)* back-transformation of the segmented labels into the original space; *(f)* post-processing to identify the valid cones. The details of each step are described below. The mathematical explanation of each step is given in supplementary.

#### (a) Global alignment of the eye

In general, the position and orientation of the eye in the volumetric data is arbitrary, depending on how the sample was mounted during scanning. However, later steps of the analysis rely on the eye surface being approximately aligned with the *xy*-plane. Therefore, the first step of the analysis is to rotate the eye.

To align the eye, we used the previously segmented (see *Volumetric segmentation of the eye*) volumetric images (*tiffs*, *nifty*, or any other image file format), where voxels labeled as cornea formed a point cloud that represented the surface of the eye. Using Principal Component Analysis (PCA), as shown in *Figure 1a,* the eye was rotated to have the longest side (first principal component) aligned with the *x*-axis, and the external-to-internal (or proximodistal) direction (third principal component) aligned with the *z*-axis. This step allows for a more systematic procedure that facilitates comparison and automation by transforming the set of points corresponding to the label of the external surface of the cornea into a coordinate system called ‘PCA space’, (*x’, y’*, z’).

#### (b) Division into sub-regions, local alignment and surface modelling

The goal of this step is to obtain a sensible 2D coordinate system of the compound eye by building a 3D polynomial surface model for the external cornea voxels. For the eyes that have high curvature, such as the *Pieris napi* sample that has an almost hemispherical eye, it would be difficult to achieve an accurate polynomial fitting on the complete external cornea. We therefore divided the external surface of the cornea of such highly curved eyes into sub-regions in the ‘PCA space’. Practically, the division of the external surface of the cornea can be realized by defining a grid of 2D rectangular masks in the dimension of (*x’, y’*), with a small overlapping area at the borders of the sub-regions, in order to avoid the loss of cones near the borders of the sub-regions in later steps.

The cornea surface in each sub-region was then locally aligned by preforming a PCA-based rotation as in *(a)*. With the eye surface roughly aligned with the *xy*-plane, we could fit a polynomial surface *z=f(x,y)* to the voxels labelled as cornea. We chose to fit a fifth order polynomial surface. An example of the surface fitting on a sub-region of the honeybee eye is displayed in *Figure 1b*.

#### (c) Sub-volume extraction and unfolding

The purpose of this process is to extract a sub-volume of the original data volume including the crystalline cone layer in each sub-region.

As the external sub-surface of the cornea is now modelled as a polynomial function, it could be resampled to a matrix of points with a density that offers sufficient resolution and optimizes the computational time. Each sub-surface sample point was then paired with a unit direction vector that was defined as the surface normal at the sample point on the cornea and pointing towards the inside of the eye. By sampling along the unit vectors, we obtain a sub-volume aligned with the “flattened” cornea surface within a defined displacement along the cornea surface normal.

In order to extract the intensity of the query points, they are back-transformed to the original coordinate system, and the image intensity is sampled from the data volume, using tricubic interpolation. Due to the step-wise extraction of displaced cornea layers, the sub-volume is in some sense “unfolded”, as illustrated in *Figure 1c*.

#### (d) Texture based auto-segmentation of the cones

In the unfolded sub-volumes, the crystalline cones should ideally appear as ‘cylinders’ aligned with the z-axis (the displacement layers), with a predictable repeated pattern of small circular structures. The segmentation of cones is thus simplified to a two-dimensional problem.

In this paper, we chose the texture-based segmentation tool *InSegt* (Dahl et al., 2020). To train *InSegt*, a user selected a single slice from the unfolded volume. This slice should be within the crystalline cone layer. The user provided a sparse manual annotation of this slice by marking some pixels as belonging to the background class, and some pixels as belonging to the cone class. The corresponding full segmentation of the slice is presented to the user for inspection, such that labelling continues until segmentation is satisfactory (cyan and magenta overlay in *Figure 1d, movie S1*). Based on this input, a dictionary was trained and applied to all other slices in the unfolded sub-volume.

#### (e) Back-transformation

The labelled voxel coordinates were finally mapped back into the original coordinate system, using the transformation described in supplementary Equation 1. A label containing multiple raw segmented cones in the original 3D image was thus obtained (*Figure 1e*).

#### (f) Post-processing of the raw cones

The automatically segmented labels unavoidably contain pixels that are not cones. A post-processing is necessary to exclude noisy detections while keeping as many correctly segmented cones as possible *(Figure 1f)*. In order to differentiate, the automatically segmented labels before post-processing are hereby called “raw cones”. The post-processing was divided into three main steps: identification of the individual raw cones, calculation of the raw cone characteristics, stepwise elimination of noisy detections *(Figure S1)*.

i. Identification of individual raw cones A connected component analysis in the original coordinate system was performed to detect and individually label each raw cone (some of which are artefacts).
ii. Calculation of raw cone characteristics A PCA was performed on each raw cone to calculate its geometric properties and detect outliers. The coefficients, eigenvalues of the covariance matrix and estimated mean were used to calculate its *elongation axis*, *length* and *radius*, and *cone centre*, respectively. The sign of the *elongation axis* was chosen so that it points towards the surface of the cornea. The *cone size* was the number of voxels included in the raw cone volume. The *distance to neighbouring cones* was the average distance between the raw cone centre and the centres of the three closest neighbouring raw cones.
iii. Stepwise elimination of noisy detections In this part, the cone characteristics were used to detect outliers in four consecutive sorting steps:

1. Raw cones with a *cone size* lower than ten voxels were immediately removed because they were likely to be noisy detections.
2. As in an earlier step (see *Modelling of the outer cornea surface*), the coordinates of the *cone centres* were transformed in the ‘PCA space’ given by the coefficients of a PCA of the cornea surface in each eye sub-region. A polynomial fitting (method *poly55*) was implemented on the normalised transformed *cone centres* using the *z*-component as a dependent variable. In each eye sub-region, the residuals of the polynomial fitting were used to cluster the cone candidates into one, two or three groups according to the gaussian mixture model (function *fitgmdist*) with the best Akaike Information Criterion (AIC). The user then indicated through visual inspection which groups of raw cones appeared valid in each eye subregion. Invalid cones generally stick out from the clusters of valid cones that gradually follow the eye curvature, making visual inspection relatively fast (~2 min per specimen) and straightforward. Groups of raw cones that were not validated by the user were discarded in the following steps.
3. The coordinates of the remaining *cone centres* were transformed into the coordinate system now given by the coefficients of a PCA of the full cornea surface. Five polynomial (*poly55*) fittings were implemented in the (*x, y*) components of the *cone centres* on the following dependent variables: normalised *z*-component of the *cone centres, size*, *length*, *radius* and *distance to neighbouring cones*. These variables were chosen for the detection of outliers because they are conserved in the true cones (expect from small topological variations) but stand out in the noisy detections. The detailed procedure for the detection of outlier is provided in *Text S1*.
4. A final polynomial fitting (*poly55*) was implemented on the *cone centres* of the remaining raw cones. A linkage was computed to determine the proximity of the fitted *cone centres* onto the polynomial surface. This linkage was used to define clusters of fitted *cone centres* that were closer than the *cut-off distance*. The *cut-off distance* was set by the user at 13 μm for all specimens because the average distance between ommatidia is approximately 20 μm in these specimens, such that two cones separated by less than a *cut-off distance* are abnormally close. In each cluster of fitted *cone centres*, the raw cone that had the largest fitting residual was eliminated iteratively until the distance between all the fitted *cone centres* in the cluster was greater than the *cut-off distance*.

The raw cones that had not been disqualified throughout the four elimination steps were classified as valid cones.

### Performance of the auto-segmentation process

To assess the performance of *InSegtCone*, five metrics were calculated. The surface modelling error and angular discrepancy between automatic and manual segmentation reflect the accuracy of the method (using manual segmentation as a reference), whereas the segmentation time, percentage of auto-segmented cones and local cone density ratio of cones reflect the efficiency of the method.

#### (a) Surface modelling error

During surface modelling (see *division into subregions and surface fitting*), the *surface modelling error* was calculated as the average of the root mean squared error (rmse) of the polynomial fittings of the surface of the cornea across each of the eye sub-regions.

#### (b) Segmentation time of the auto-segmentation method

To evaluate the performance of the auto-segmentation method, the segmentation time of the process was estimated. The volumetric segmentation of the eye is probably the most time-consuming step of the analysis (2-4 hours per specimen), but was not taken into account here because it is a prerequisite but not a part of the auto-segmentation of the cones. The segmentation time of the auto-segmentation process was thus approximately equal to the longest step – the texture-based auto-segmentation. When evaluating the texture-based segmentation duration, we considered only the cumulated time spent by the user to manually annotate and train the segmentation model on each eye sub-region. The rest of the computing time for automatic segmentation using the trained dictionary highly depends on the size of the image data and the computer power, therefore it is not discussed in detail here.

#### (c) Percentage of auto-segmented cones

This was the number of valid cones after post-processing divided by the predicted number of ommatidia that was obtained using the method described in Taylor et al. (2019a). In brief, manual measurements of facet diameters at about 30 locations across the eye were used to build an interpolant of facet area across the eye surface. The predicted number of ommatidia is thus the total area of the cornea surface estimated from an *isosurface fit* (function *isosurface* with *isovalue* = 0.5) divided by the average interpolated facet area.

#### (d) Angular discrepancy between automatic and manual segmentation

The same cone can be segmented both manually in Amira and with the auto-segmentation method, thus generating two *cone duplicates*. This was considered to be the case if the *cone centre* of an auto-segmented cone was less than a voxel away from a manually segmented cone. The angular discrepancy between automatic and manual segmentation was the angle *α ∈ [0, π]*, between the elongation axes of the two *cone duplicates*.

#### (e) Local cone density ratio

The local cone density ratio *R ∈ [0, 1]*, represents the local coverage of the auto-segmentation method. A ratio equal to 1 indicates that all cones predicted locally were identified and segmented by the method. To calculate *R*, equidistant sampling points on the cornea surface were obtained as in Taylor et al. (2019a) . The local density ratio at each sampling point was the number of cones identified by the auto-segmentation method within a given range (100 μm) divided by the expected number of ommatidia in the same range. The latter was the local area of the cornea subset divided by the local average facet area.

## Results

When applied to compound eyes of *Apis mellifera*, *Bombus terrestris* and *Pieris napi*, *InSegtCone* enabled the automatic reconstruction of 4423, 3783 and 4334 cones, corresponding to 80%, 76% and 62% of the predicted total number of crystalline cones, respectively (*figure 2* and *figure 3c, movies S2-4*). The cone density ratio *R*, i.e. the local ratio of auto-segmented over predicted cones, was close to 1 across most regions of the eyes of *A. mellifera* and *B. terrestris*, indicating that all the cones predicted in these areas were segmented (*figure 2*). The value of *R* dropped in the most dorsal region and at the edges of the eye, indicating that the cones that remained undetected were situated in these areas (representing ~20% of the total number of cones). The topography of *R* was more irregular in *Pieris napi*, revealing small areas where cones were not segmented, in the most dorsal and ventral regions and at the edges of the compound eye.

**Figure 2:**
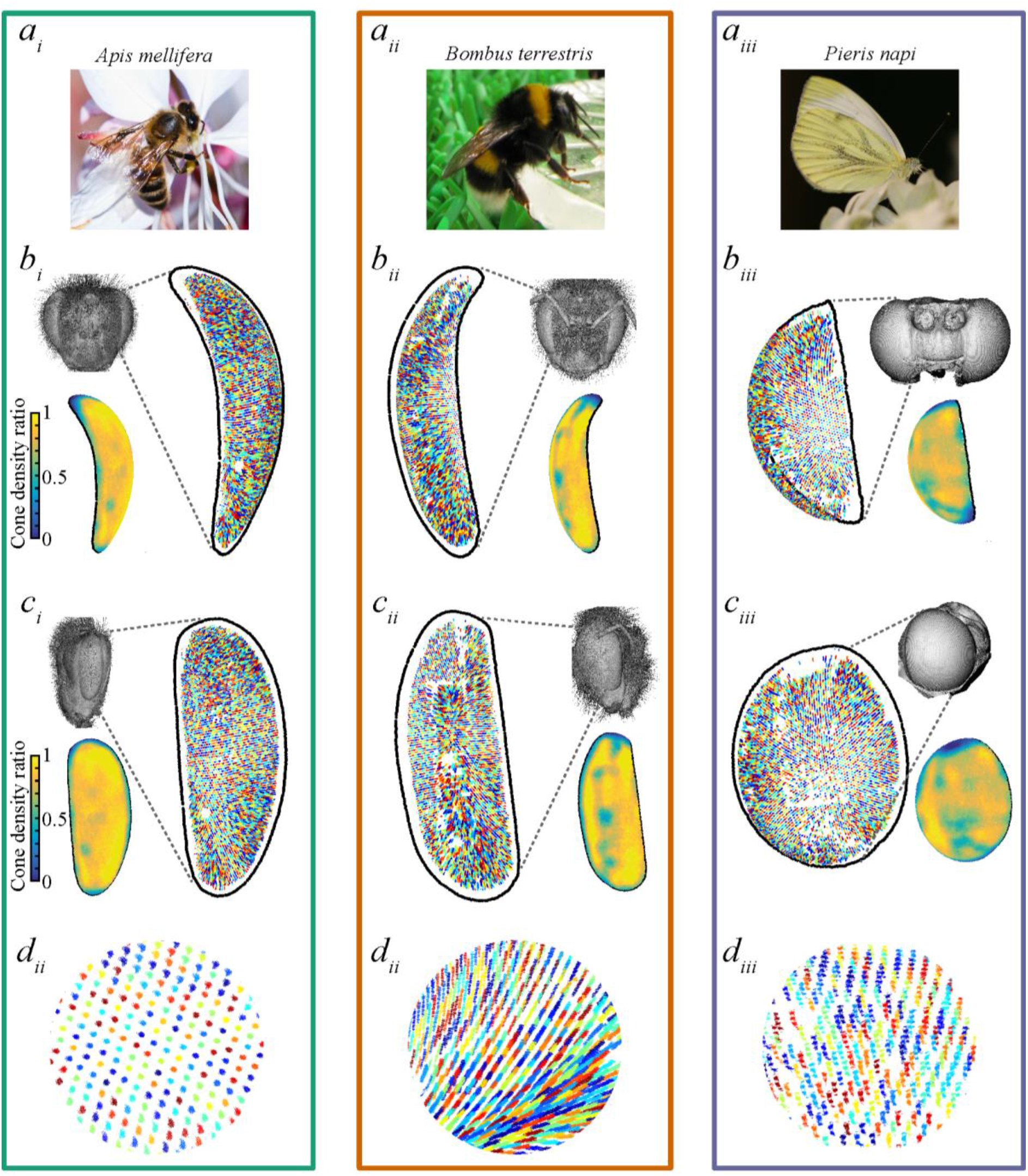
Auto-segmented crystalline cones in the compound eyes of three arthropods. The eyes of *Apis mellifera* (*a_i_*), *Bombus terrestris* (*a_ii_*) and *Pieris napi* (*a_iii_*) were divided respectively into 4, 12 and 9 subregions. Overview of the cones segmented with *InSegtCone* (each labelled with a different colour) from the front *(b_i-iii_)* and the side *(c_i-iii_)*, as indicated by the volume renderings of the species’ heads (grey images). The border of the external surface of the cornea is represented in black. The miniature in each left corner indicates the topology of the cone density ratio. A ratio locally equal to one means that all expected cones were auto-segmented. *(d_i-iii_)* Detailed view of a portion of the auto-segmented cones.

**Figure 3:**
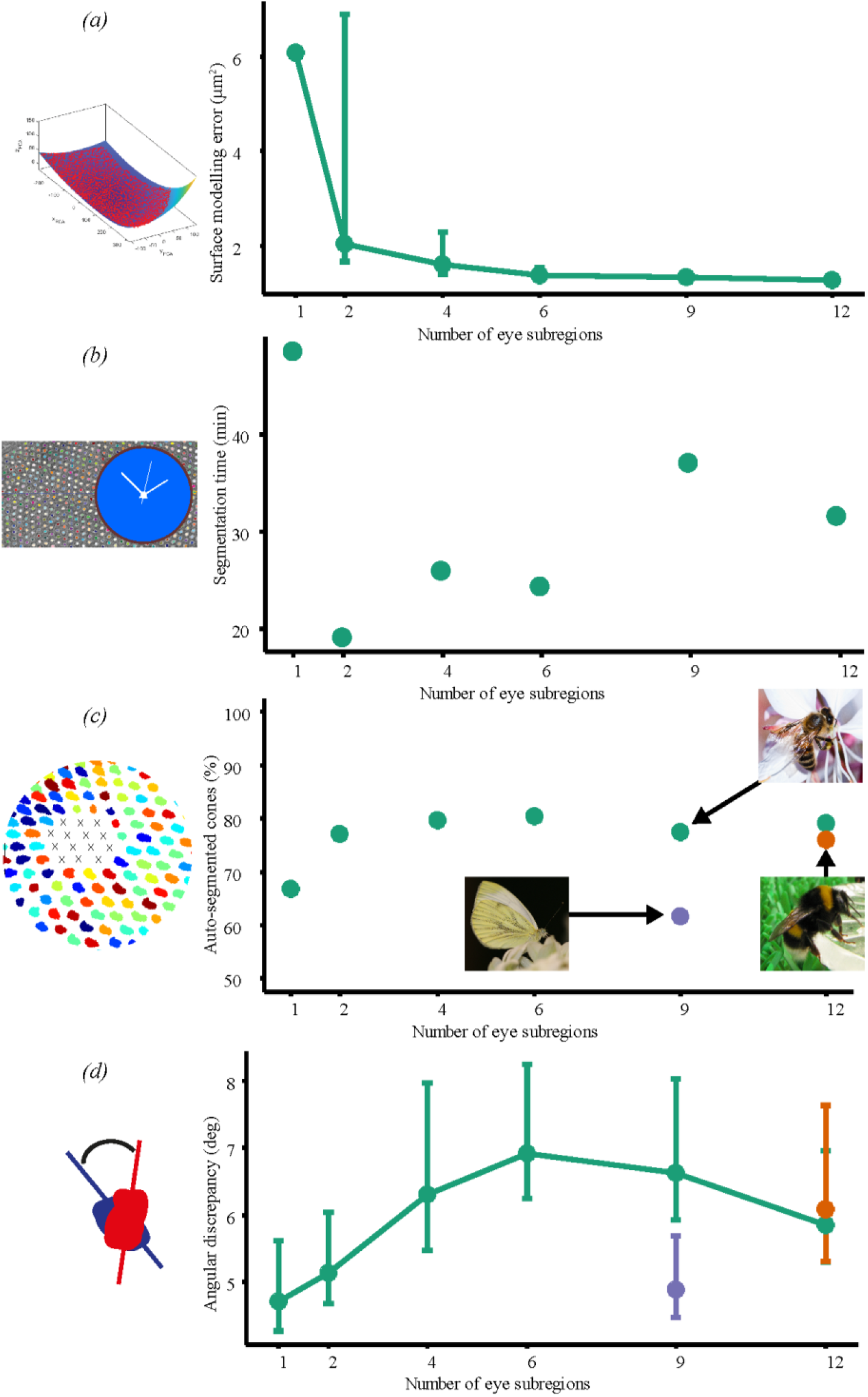
Performance of the auto-segmentation method in *Apis mellifera* (green), *Bombus terrestris* (orange) and *Pieris napi* (purple). The surface modelling error *(a)*, segmentation time *(b)*, percentage of auto-segmented cones *(c)* and angular discrepancy between manual and auto-segmentation *(d)*, are represented as a function of the number of eye subregions used in the segmentation algorithm. When drawn, error bars represent the standard error around the mean values.

Once the segmentation of the eye and cornea (that requires most manual work) were completed, the segmentation time of the automatic method was ~30 min for ~4400 cones (*figure 3b*), whereas the duration of the manual segmentation was ~180 min for 100 cones (table S1). This means that the manual method permitted the segmentation of one cone in about 2 min when the automatic method segmented one cone in less than 0.5 s.

To test the effects of dividing the eye into sub-regions, the auto-segmentation process was applied to the compound eye of *A. mellifera* when it had been divided into increasing numbers of sub-regions *NSR*. The aim of this test was also to identify the optimal *NSR* that trades off accuracy and efficiency. The percentage of segmented cones initially increased with increasing *NSR* (*NSR ∈ [1, 4]*, *figure 3c*), until it reached a plateau and even slightly dropped (*NSR ∈ [4, 12]*, *figure 3c*). This initial gain highlights the benefits of dividing the eye into sub-regions, and was due to the segmentation of additional cones in the dorsal area of the compound eye (*figure S2*). This was likely facilitated by the initial decrease of the surface modelling error (*figure 3a*). The improved efficiency of the algorithm up to *NSR* = 4 did not increase the run-time of the auto-segmentation process (*figure 3b*), although it may have caused a slight increase in the average angular discrepancy between the automatic and the manual segmentation (*figure 3d*).

The angular discrepancy *α* between automatic and manual segmentation, that is the angle between the elongation axes of the manually and the auto-segmented duplicates of the same cone, could locally be close to 70 deg (*figure S4*). Note that this does not indicate that the auto-segmented cone is 70 deg off the elongation axis, as manual segmentation is also prone to errors. Moreover, *α* was on average below 7 deg for all eyes (*figure 3d*), which indicates a good agreement between the manual and the auto-segmentation methods overall.

## Conclusion

In this paper, we demonstrate *InSegtCone,* a new computational method for automatically segmenting the crystalline cones of compound eyes in arthropods using high-resolution micro-CT images. We assessed the performance of the method by implementing the auto-segmentation process on the apposition compound eyes of three insect species. Our method labelled most of the crystalline cones across of the compound eyes and generated cone labels with a similar level of accuracy than manual segmentation but ~250 times faster. We conclude that this new method for auto-segmentation of crystalline cones is accurate and efficient. *InSegtCone* sets the ground for subsequent high-throughput analyses that are required for understanding the diversity of eyes and vision in arthropods.

### Current limitations and possibilities for improvement

Because the present automatic procedure is ~250 times faster than manual labelling, it greatly facilitates the segmentation of crystalline cones and minimises most of the labour-intensive manipulations usually required for such analyses. However, the method still requires some level of manual interaction with the user, in particular, during the segmentation of the external surface of the cornea and the texture-based segmentation that requires the annotation of crystalline cones on a 2D slice (if no existing dictionary from similar samples is available).

There are thus opportunities for further automation of the segmentation process. We encourage future users to carefully consider the advantages of using the auto-segmentation instead of a manual segmentation method. The automatic method is probably most beneficial when applied to numerous eye samples and/or species with many ommatidia (and thus many cones), such as the bees and butterfly presented in this paper that possess several thousands of ommatidia. Manual segmentation may be more advantageous for studies restricted to few specimens of arthropods with a small number (<100) of ommatidia (Taylor et al., 2020).

Our auto-segmentation tool uses clustering of image features and manual labelling from one slice to segment repeated patterns (here, in the form of small circles). It will perform well on images that have a similar appearance as the image used for training the dictionary. However, micro-CT reconstructions sometimes generate artefacts (e.g. ring artefacts, beam hardening, etc.) that affect the appearance of the cones and can locally disrupt the auto-segmenting of cones. An example of this is indicated by the arrow in *figure 1d* where the algorithm failed to segment the cones around the stripes caused by ring artefacts. Fortunately, micro-CT is a non-destructive imaging method that allows repeated scanning on the same specimen with adjusted parameters to enhance image quality and limit artefacts. The image quality can also be improved during reconstruction with numerous post-processing tools, such as ring removal algorithms (Vo et al., 2018).

During the texture-based segmentation step, a *training dictionary* is built from the manual annotation of a training slice (Dahl et al., 2020). If the cross sections of the cones have a highly variable appearance across the slice, the use of a unique training dictionary common to all the cones is likely to lead to inaccurate or poor segmentation results. The variable appearance of cones across a training slice can have several explanations. For example, *(1)* if the thickness of the cornea significantly changes across the eye, the training slice will contain cones that appear different because they are viewed at different positions along their main elongation axes. This issue may be avoided by using a more robust reference for surface modelling, such as the internal, instead of the external, corneal surface, although this would require additional volumetric segmentation efforts. Another solution is to divide the compound eye into a higher number of sub-regions to ensure that the thickness of the cornea is homogeneous within each sub-region. *(2)* The cones themselves can have distinct morphologies in different regions. For instance, the cones located in the dorsal area of the bee eyes appear to be shorter and more densely packed than in the rest of the eye (*figure 2*). In this case, a sub-region dedicated to these challenging regions may be needed to improve segmentation. *(3)* Auto-segmentation can be complicated if neighbouring crystalline cones are in contact with each other, such as in the case of crab compound eyes (Alkaladi and Zeil, 2014; Bagheri et al., 2020). In this case, the texture-based segmentation algorithm either fails, or the cones are segmented together in a bundle of connected voxels and cannot be isolated without additional post-processing, e.g. morphological erosion and dilation techniques (Gonzalez et al., 2004). An alternative solution is to combine the texture-based segmentation with shape recognition algorithms, such as circle detection (Gonzalez et al., 2004).

### Current and future Applications

Despite the limitations discussed above, *InSegtCone* represents a formidable opportunity for the study of the visual biology of arthropods. Because it greatly accelerates time-consuming labelling, this new tool enables comprehensive studies across a large number of arthropod eyes, which had been practically inaccessible until now. The auto-segmentation method also has a wide range of potential applications within and beyond the study of the anatomy of compound eyes. Firstly, (and possibly most obviously) the labels of the crystalline cones can serve to generate unbiased functional optical models of the eyes. This is because, without analysis of the crystalline cones, optical models must be based on the apparent angular spacing of facets on the external surface of the cornea, the corneal IO angle, which produces biased estimates of optical resolution and of the field of view (FOV) of the eye (Bergman and Rutowski, 2016; Taylor et al., 2019a). The auto-segmented labels can be used to estimate the skew *β* between the normals to the external cornea and the main *elongation axes* of the crystalline cones across the compound eye (Stavenga, 1979). The cone skew *β* is then necessary to compute the visual IO angle – an unbiased estimate of optical resolution– that can be mapped across the accurately delineated FOV of the eye. Ultimately, the fast auto-segmentation method makes it possible to compare theses accurate resolution maps across large numbers of specimens with different sizes, life-stages, sex, species, etc. This is likely to advance our understanding about the visual ecology and evolution of vision in arthropods (Bagheri et al., 2020; Scales and Butler, 2016; Somanathan et al., 2009; Taylor et al., 2019a). The auto-segmentation method not only extracts the *elongation axis* of the crystalline cones but also their full shape. This is interesting because several arthropod species modify the length (Brodrick et al., 2020; Menzi, 1987; Nilsson and Odselius, 1981) and the diameter (Brodrick et al., 2020) of their crystalline cones in response to changes in light levels. In fiddler crabs, these light-adaptation mechanisms together with modifications of the rhabdoms enhance optical sensitivity at night (Brodrick et al., 2020). The auto-segmentation method generates opportunities for large scale investigation of these light-adaptation properties at numerous light levels and across large numbers of species.

Applying the auto-segmentation method to other parts of the eyes could facilitate the labelling of other elements in the compound eyes. For instance, the centre of individual facets could be identified automatically by running the program superficially over the external surface of the cornea. This technique would promptly generate 3D maps of facet dimensions across the eye and thus good estimates of the total number of ommatidia. The program could be adapted to extract morphological properties of the rhabdoms when they are visible, i.e. when the resolution and contrast of micro-CT images are sufficient. This would represent a great benefit given that current studies of rhabdom morphology require tedious sample preparation and slicing for Transmission Electron Microscopy. A modified version of the auto-segmentation method could be used to measure the diameter of rhabdom, either by using the proximal tip of the segmented crystalline cone, or directly by segmenting the rhabdom shape. Besides being more accurate, the latter method would have the additional advantage of extracting other properties, such as the length of the rhabdoms. To segment rhabdoms, we expect better results using the retina-cone interface as a reference during the surface modelling step, rather than the external surface of the cornea. With these measurements of rhabdom diameters and lengths, scientists would be able to calculate the topology of optical sensitivity with unprecedent level of detail across the eye.

Finally, *InSegtCone* could solve segmentation problems beyond the study of compound eyes. In micro-CT scans of camera-type eyes, such as ocelli (Wilby et al., 2019), this represents a promising tool for fast reconstruction of the shape of photoreceptors across the retina. In principle, our method can be extended to label photoreceptors on any tomographic reconstruction of vertebrate or invertebrate retina, regardless of the imaging technique involved to generate data (micro-CT, confocal microscopy, etc). More generally, this work can inspire projects in a wide range of fields that require tools to segment repeated elements in a 3D layer. In Biology, these repeated elements are for example: muscle fibres, olfactory sensilla on antennas, epidermal appendages such as scales, vascular tissue and roots of plants. The first part of the process involving the modelling of the external cornea may also inspire other studies to achieve better visualization or easier quantitative analysis through the unfolding of 3D images.

## Acknowledgments

We would like to thank Carina Rasmussen, Eva Landgren, Ola Gustafsson, Karin Odlén, Julia Källberg, Per Alftrén, Viktor Håkansson, Zahra Moranidour, Craig Perl, Vun Wen Jie, Marie Guiraud, Zanna Johansen, Gavin Taylor and Marie Schmidt for help during sample preparation and scanning. Thanks to Qiang Tao, David Wilby, Rajmund Mokso, Andrew Bodey, Christoph Rau and Kazimir Wanelik for their assistance during imaging at Diamond Light Source (proposals 13848 and 16052). We thank Gavin Taylor, Marie Schmidt, Valentin Gilet and Viktor Håkansson for help during data analysis. Emily Baird is thankful to the Swedish Research Council (201806238 and 2014-4762) and the Lund University Natural Sciences Faculty for financial support. Pierre Tichit was supported by a grant from the Interreg Project LU-011. Data acquisition at SUBIC was supported by grant SU FV-5.1.2-1035-15. This work was supported in part by The Center for Quantification of Imaging Data from MAX IV (QIM) funded by The Capital Region of Denmark.

## Data availability

The original image data of the compound eyes as well as the manual and automatic cone labels will be made available on *MorphoSource* prior to publication. The raw data used to evaluate the performance of the method will be available on *Dryad*. The MATLAB scripts used for the auto-segmentation process will be made available on *Github* upon publication.

## Author contributions

All authors designed the study. HMK, TZ and PT collected the data and generated the analyses. HMK, TZ and PT drafted the manuscript. All authors revised the manuscript.

