## Supplementary Material for "*InSegtCone*: Interactive Segmentation of crystalline Cones in compound eyes"

### *InSegtCone*: Interactive Segmentation of crystalline Cones in compound eyes Supplementary Material

### Text S1: Supplemental method of the Automated Segmentation of the crystalline Cones

The auto-segmentation process of crystalline cone consisted of six steps: (a) reorientation of the eye; (b) dividing the outer cornea voxels into subregions and modelling as polynomial surfaces; (c) extraction and unfolding of a sub-volume of the data capturing the crystalline cone layer; (d) auto-segmentation of the raw cones using a texture-based approach; (e) back-transformation of the segmented labels into the original space; (f) post-processing to identify the valid cones. The mathematical explanation of each step is given here for each step.

(a) *Reorientation of the eye*

The reorientation of the eye is achieved through a Principal Component Analysis (PCA) (Stegmann 2002), which simplifies the process of estimating a surface of the cornea, as described here:

Let the coordinates of all cornea-labelled voxels, $\mathbf{p}_i$, be stored as a $N_p$×3 point set, $\mathbf{P} = \{\mathbf{x_p}, \mathbf{y_p}, \mathbf{z_p}\}$. A PCA of $\mathbf{P}$ is then carried out, considering each voxel as an observation with 3 variables. The analysis provides an estimate of the point set centroid, $\mathbf{p}_\mathrm{C}$ (the column-wise mean of $\mathbf{P}$), and the 3×3 matrix of the orthogonal eigenvectors, $\mathbf{V}$.

Using those, the cornea-labelled voxels $\mathbf{p}_i$ are transformed into a coordinate system called 'PCA space', $(x', y', z')$, where it has the smallest possible bounding box,

$$\mathbf{p}_i' = \mathbf{V}(\mathbf{p}_i - \mathbf{p}_\mathrm{C}), i = 1, \ldots, Np \qquad \textit{Equation 1}$$

The PCA transformation represents an unbiased way of removing translational and rotational sources of variation, caused by for example the sample placement and orientation in the scanner.

*(b) Division into subregions and surface modelling*

The goal of this step is to obtain a 2D coordinate system of the compound eye by building a 3D surface model for the external cornea voxels.

For eyes with high curvatures we divide the external surface of cornea into subregions in the 'PCA space'. For convenience, the subregions are defined as rectangles in a grid shape that covers the area of cornea surface. The cornea point set $\mathbf{P}_m' = \{\mathbf{x}_m', \mathbf{y}_m', \mathbf{z}_m'\}$ in the $m^{\mathrm{th}}$ subregion ($m$ = 1, …, $N_{\mathrm{SR}}$) is then once again transformed into a new 'PCA space' ($x_m'', y_m'', z_m''$) following the same method as $\mathbf{p}i' = \mathbf{V}(\mathbf{p}_i - \mathbf{p}_\mathrm{C}), i = 1, \ldots, Np$ *Equation 1*,

$$\mathbf{p}_{i,m}'' = \mathbf{V}_m'\left(\mathbf{p}_{i,m}' - \mathbf{p}_{\mathrm{C},m}'\right), i = 1, \ldots, N_{p_m},$$

where $\mathbf{p}_{\mathrm{C},m}'$ is the point set centroid for the points in the m$^{th}$ subregion and $\mathbf{V}_m'$ the 3×3 matrix of the orthogonal eigenvectors.

Finally, a 5$^{\mathrm{th}}$ order polynomial surface is fitted to the cornea point set in each subregion $\mathbf{P}_m'' = \{\mathbf{x}_m'', \mathbf{y}_m'', \mathbf{z}_m''\}$. Let $\boldsymbol{\beta}$ be the vector of coefficients for the polynomial function, *f*. An estimate, $\widehat{\boldsymbol{\beta}}$, can be obtained by solving the least squares problem,

$$\widehat{\boldsymbol{\beta}_m} = \min_{\beta_m} \| f(\mathbf{x}_m'', \mathbf{y}_m'', \boldsymbol{\beta}_m) - \mathbf{z}_m'' \|_2^2$$

Note, that the 1st and 2nd principal axis coordinates, $\mathbf{x}_m''$ and $\mathbf{y}_m''$, are used as observations of the independent variables and the 3rd axis coordinates, $\mathbf{z}_m''$, as the dependent variable. Without the transformation into the 'PCA space', the selection of variables for the fitting would not be automatically given.

*(c)* *Sub-volume extraction and unfolding*

The purpose of this process is to extract a $Nx \times Ny \times N_l$ sub-volume, $\mathbb{Q}_m$, of the original data volume including the crystalline cone layer in the subregion $m$ ($m = 1, \ldots, N_{SR}$).

The external subsurface of the cornea is now described as a polynomial function, $f_m(x''_m, y''_m, \widehat{\boldsymbol{\beta}_m})$, $x''_m \in [a_x, b_x]$, $y''_m \in [a_y, b_y]$, and it could hence be resampled to any desired density using respectively $Nx$ and $Ny$ points in the defined function intervals. Each subsurface sample point, $\mathbf{s}_{(i,j)} = \{i, j, f_m(i, j, \widehat{\boldsymbol{\beta}_m})\}$, was then paired with a unit direction vector, $\mathbf{d}_{(i,j)}$, that is defined as the surface normal at the sample point on the cornea and pointing towards the inside of the eye. Let $\Delta_D$ represent a displacement length and $N_l$ a number of steps. This forms the basis for defining at set of query points

$$\mathbf{q}''_{(i,j,n)} = \mathbf{s}_{(i,j)} + (n-1) \cdot \Delta_D \cdot \mathbf{d}_{(i,j)},$$
$$i = 1, \ldots, N_x, \quad j = 1, \ldots, N_y, \quad n = 1, \ldots, N_l$$

where ($i$, $j$) represents a coordinate on the cornea surface and $n$ represents a displacement layer.

The total set of query points are then back-transformed to the original coordinate system, and the image intensity is sampled from the data volume, $\mathbb{D}$, using tricubic interpolation

$$\mathbb{Q}_m(i,j,n) = \mathbb{D}\left[\left(\boldsymbol{q}''_{(i,j,n)} \mathbf{V}'^{\,T}_m + \mathbf{p}'_{C,m}\right)\mathbf{V}^T + \mathbf{p}_C\right], \qquad \textit{Equation 2}$$
$$i = 1, \ldots, N_x, \; j = 1, \ldots, N_y, \; n = 1, \ldots, N_l.$$

*(d)* *Texture-based auto-segmentation of the raw cones*

The detailed explanation of the texture-based segmentation tool *InSegt* can be found in the original publication (Dahl et al., 2020).

*(e)* *Back-transformation*

The labelled voxel coordinates could finally be mapped back to the original coordinate system, using the transformation described in *Equation 2*.

*(f)* *Post-processing of the raw cones*

iii. Stepwise elimination of noisy detections

*i.* Five polynomial (poly55) fittings were implemented in the ($x$, $y$) components of the *cone centres* on the following dependent variables, $v$: normalised $z$-component of the *cone centres*, *size*, *length*, *radius* and *distance to neighbouring*. Let $r_i$ be the residual of the fit on the variable $v$ at the coordinates ($x_i$, $y_i$) for the raw cone $i$. To detect cone outliers, the median absolute deviation (MAD) was calculated:

$$\mathrm{MAD(v)} = -\frac{1}{\sqrt{2} \cdot \mathrm{erfc}^{-1}\left(\frac{3}{2}\right)} \cdot \mathrm{median}(|\mathrm{r_i} - \mathrm{median(r)}|)$$

where $\mathrm{erfc}^{-1}$ is the Inverse Complementary Error Function, which can be realized by the MATLAB function erfcinv.

For each variable $v$ and raw cone $i$, the sorting criterium $SC$ was:

$$SC(v,i) = |r_i - \mathrm{median}(r)| - SP \cdot \mathrm{MAD}(v)$$

where $SP$ was the *sorting power* set by the user at three for the bee specimens, and at five for *P.napi* because visual inspection revealed that it captured most true cones and discarded all noisy detections.

For every variable *v*, a raw cone *i* was considered as an outlier and discarded if *SC* was greater or equal to zero. Essentially, a raw cone whose fitting residual deviated of more than *SP* times the MAD from the median of the fitting residuals was eliminated.

### Table S1

| *Species* | *Facet diameter (µm)* | *External lens surface (µm²)* | *Number of subregions* | *Surface modelling error (rmse)* | *Predicted number of ommatidia* | *Number of auto-segmented cones* | *Number of manually segmented cones* | *Auto-segmentation time (seconds)* | *Number of manual and automatic duplicates* |
|---|---|---|---|---|---|---|---|---|---|
| *Pieris napi* | 20.7±1.2 | 259130 | 9 | - | 7031 | 4334 | 104 | 2852 | 65 |
| *Bombus terrestris* | 22.8±1.9 | 224420 | 12 | - | 4977 | 3783 | 102 | 3660 | 81 |
| *Apis mellifera* | 22.6±1.9 | 244050 | 4 | 1.61±0.37 | 5548 | 4423 | 101 | 1572 | 94 |
| *Apis mellifera* | 22.6±1.9 | 244050 | 1 | 6.08 | 5548 | 4423 | 101 | 2910 | 74 |
| *Apis mellifera* | 22.6±1.9 | 244050 | 2 | 2.04 | 5548 | 4423 | 101 | 1165 | 88 |
| *Apis mellifera* | 22.6±1.9 | 244050 | 6 | 1.37±0.17 | 5548 | 4423 | 101 | 1477 | 95 |
| *Apis mellifera* | 22.6±1.9 | 244050 | 9 | 1.34±0.14 | 5548 | 4423 | 101 | 2230 | 91 |
| *Apis mellifera* | 22.6±1.9 | 244050 | 12 | 1.28±0.08 | 5548 | 4423 | 101 | 1907 | 94 |

*Table S1:* Values used for the evaluation of the performance of the auto-segmentation method. The full list of angular discrepancies between the manually and auto-segmented cones can be downloaded (as .csv files) from Dryad.

### Movies S1-S4

*Movie S1:* Texture based segmentation of crystalline cones in *Bombus terrestris* using the tool *Insegt.*

*Movie S2:* Auto-segmented crystalline cones in *Apis mellifera.*

*Movie S3:* Auto-segmented crystalline cones in *Bombus terrestris.*

*Movie S4:* Auto-segmented crystalline cones in *Pieris napi.*

### Figures S1-S4

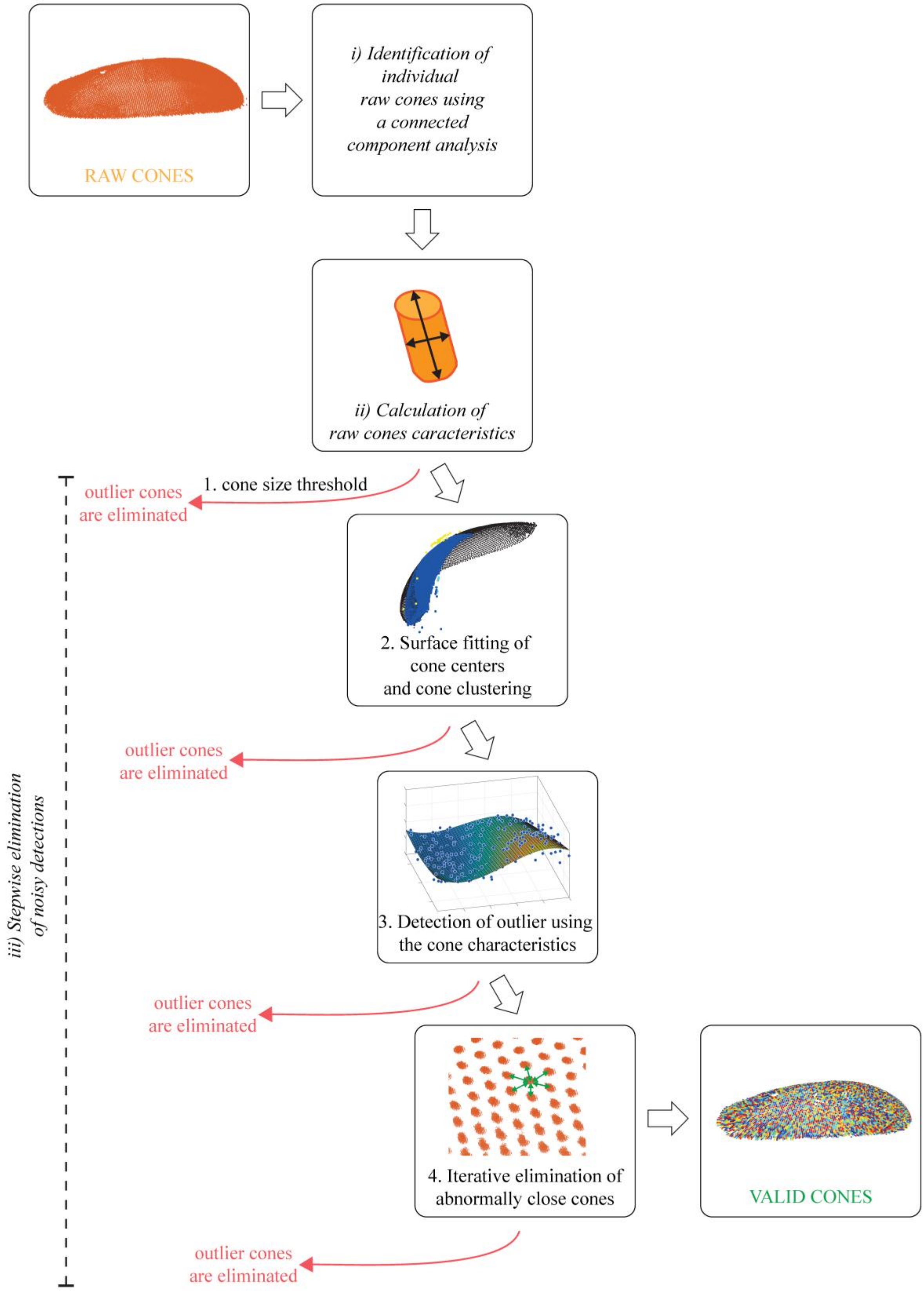


*Figure S1:* Outline of the post-processing method. The goal of this step is to sort the cones in order to distinguish between valid cones and noisy detections.

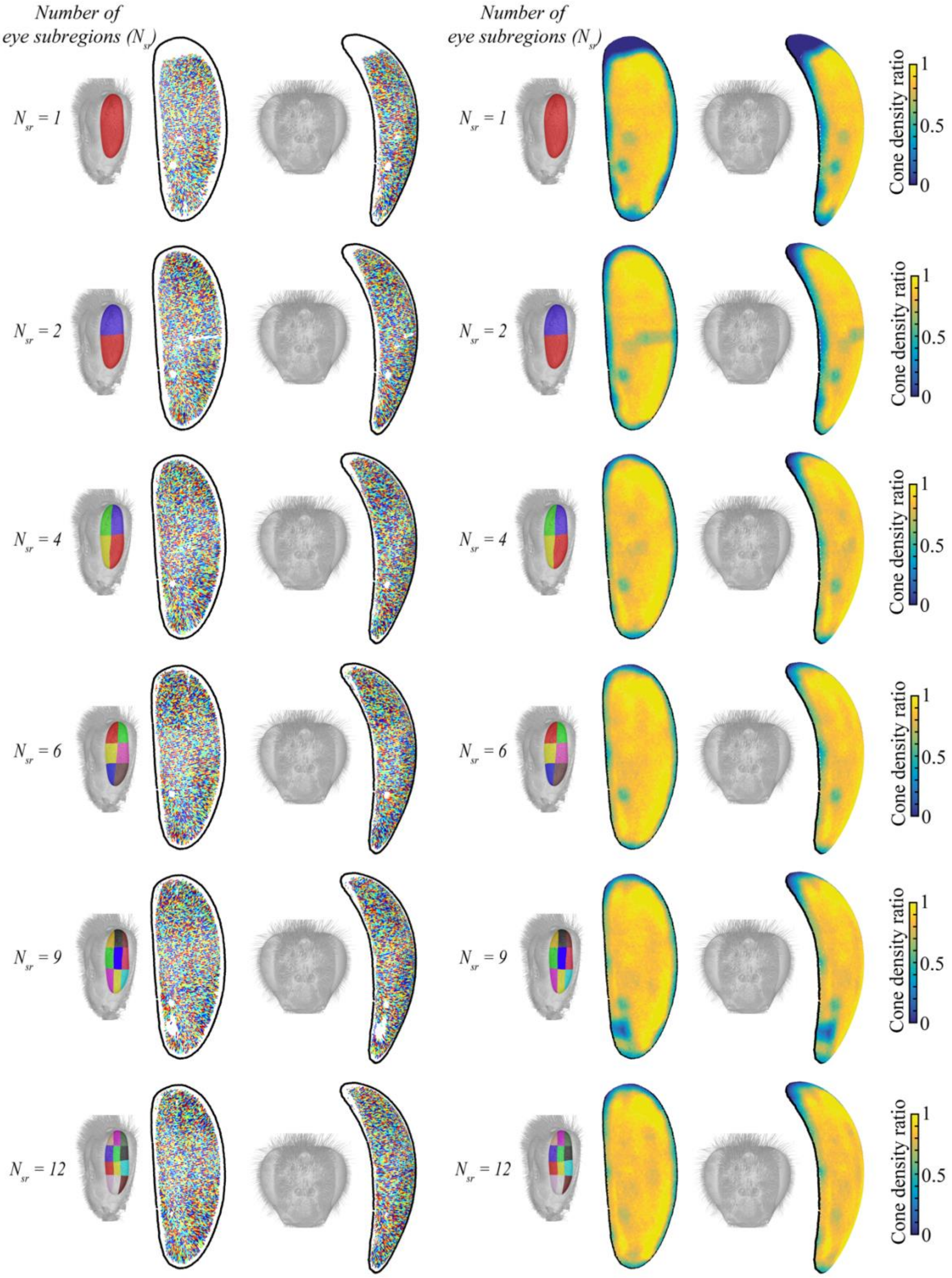


*Figure S2:* Effect on the auto-segmented crystalline cones of dividing the compound eye into subregions in *Apis mellifera*. The compound eye was divided into increasing numbers of subregions $N_{sr}$ from 1 to 12. *Left*: Overview of the segmented cones (each labelled with a different colour) from the front and the side as indicated by the volume renderings of the honeybee head. The border of the external surface of the cornea is represented in black. *Right*: topology of the cone density ratio viewed from the front and the side. A ratio locally equal to one means that all expected cones were auto-segmented.

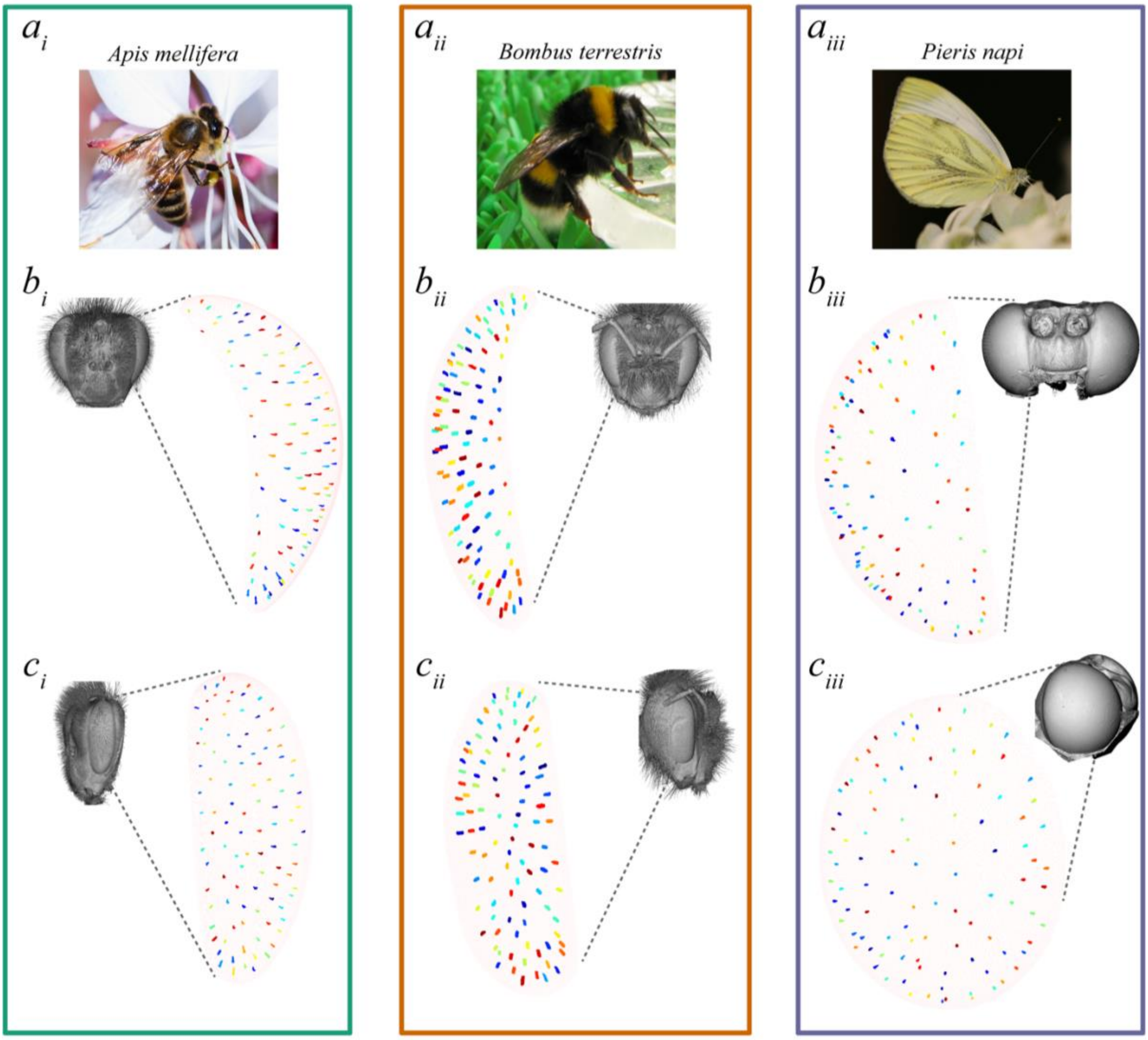


*Figure S3:* Manually segmented crystalline cones in the compound eyes of *Apis mellifera* ($a_i$), *Bombus terrestris* ($a_{ii}$) and *Pieris napi* ($a_{iii}$). Overview of the manually segmented cones (each labelled with a different colour) from the front ($b_{i-iii}$) and the side ($c_{i-iii}$), as indicated by the volume renderings of the species' heads (grey images). The border of the external surface of the cornea is represented in black.

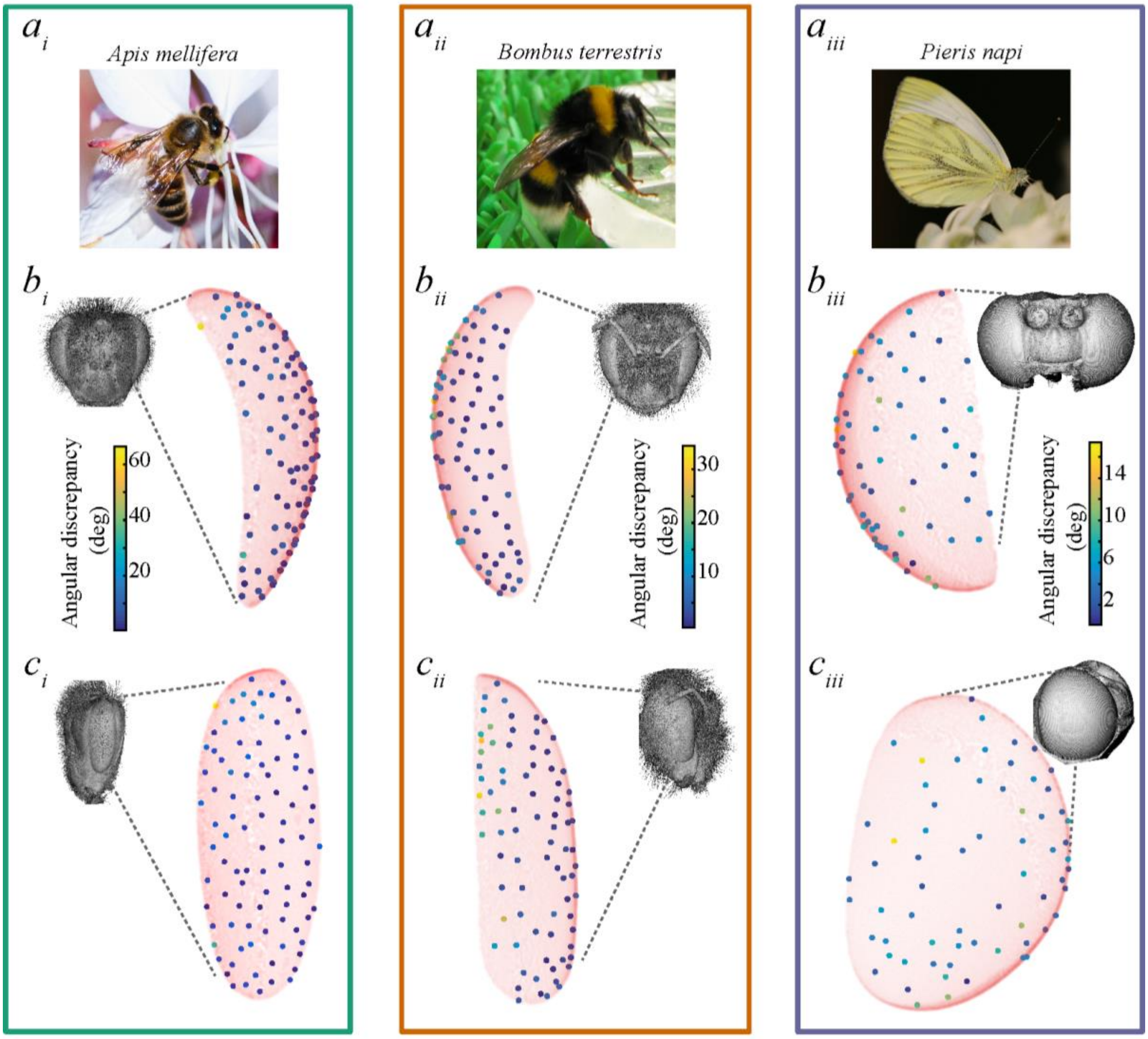


*Figure S4*: Angular discrepancies between manually and auto-segmented cones in *Apis mellifera* ($a_i$), *Bombus terrestris* ($a_{ii}$) and *Pieris napi* ($a_{iii}$). Each point corresponds to the projection of one cone duplicate on the external surface of the cornea (light pink) viewed from the front ($b_{i-iii}$) and the side ($c_{i-iii}$), as indicated by the volume renderings of the species' heads (grey images).
