## Supplementary figures and images for "*InSegtCone*: Interactive Segmentation of crystalline Cones in compound eyes"

### MovieS1

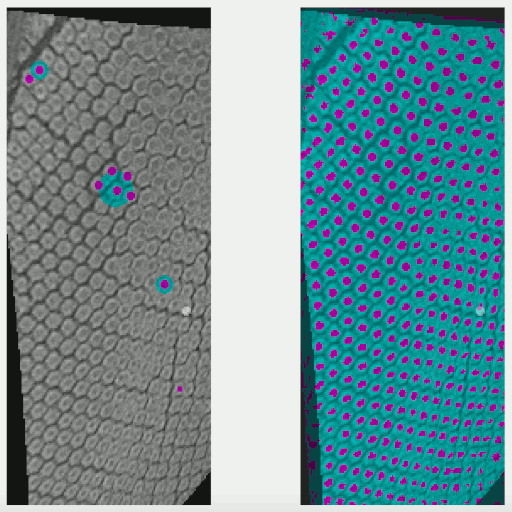
